# MutLα suppresses error-prone DNA mismatch repair and preferentially protects noncoding DNA from mutations

**DOI:** 10.1101/2024.04.01.587563

**Authors:** Lyudmila Y. Kadyrova, Piotr A. Mieczkowski, Farid A. Kadyrov

## Abstract

The DNA mismatch repair (MMR) system promotes genome stability and protects humans from certain types of cancer. Its primary function is the correction of DNA polymerase errors. MutLα is an important eukaryotic MMR factor. We have examined the contributions of MutLα to maintaining genome stability. We show here that loss of MutLα in yeast increases the genome-wide mutation rate by ∼130-fold and generates a genome-wide mutation spectrum that consists of small indels and base substitutions. We also show that loss of yeast MutLα leads to error-prone MMR that produces T>C base substitutions in 5’-A<u>T</u>A-3’ sequences. In agreement with this finding, our examination of human whole genome DNA sequencing data has revealed that loss of MutLα in induced pluripotent stem cells triggers error-prone MMR that leads to the formation of T>C mutations in 5’-N<u>T</u>N-3’ sequences. Our further analysis has shown that MutLα-independent MMR plays a role in suppressing base substitutions in N_3_ homopolymeric runs. In addition, we describe that MutLα preferentially defends noncoding DNA from mutations. Our study defines the contributions of MutLα-dependent and independent mechanisms to genome-wide MMR.

## INTRODUCTION

DNA damage and replication errors constantly challenge the integrity of genetic information. To deal with DNA damage and replication errors, the cell evolved DNA repair. The key DNA repair pathways that have been identified are homology-directed double-strand break repair, base excision repair, mismatch repair, non-homologous end joining, nucleotide excision repair, repair of DNA interstrand crosslinks, and ribonucleotide excision repair (1–7). If left unrepaired, DNA damage and replication errors cause mutations. Mutations are the molecular basis of many diseases including cancer. In addition to being detrimental, mutations are sometimes beneficial for the organism. For example, mutations that are produced in B cells during somatic hypermutation of immunoglobulin genes lead to antibody diversification, an essential step in the immune response (8).

The DNA mismatch repair system is a major DNA repair system that has been conserved from bacteria to humans (1,9). It promotes genetic stability by strand-specific removal of base-base mismatches and small insertion/deletion loops that are formed during DNA replication and strand exchange in homologous recombination (10,11). The MMR system also stabilizes genetic information by suppressing homeologous recombination and initiating apoptosis in response to irreparable DNA damage caused by several anticancer drugs (12,13). Although MMR mostly occurs in an error-free manner, it is known to be error-prone at certain genomic loci (14). The error-prone MMR causes triplet repeat expansions and the formation of base substitution mutations at A:T base pairs of immunoglobulin genes (15–17).

Studies in the bacterium *E. coli* led to the discovery and elucidation of the strand-specific MMR that is directed to the daughter strand by transient absence of DNA methylation at GATC sites (18–21). The MMR reaction in *E. coli* is initiated by recognition of a mismatch by MutS. After recognizing a mismatch, MutS recruits MutL to the heteroduplex DNA in an ATP-dependent reaction. The ternary MutS-MutL-heteroduplex complex activates MutH endonuclease to incise the daughter strand at a hemimethylated GATC site. A nick produced by MutH serves as a loading site for helicase II that unwinds a stretch of the mismatch-containing daughter strand in a MutS-and MutL-dependent reaction (22). The unwound strand is hydrolyzed by a single-stranded DNA exonuclease (ExoI, ExoVII, ExoX, or RecJ). After the excision step, the generated DNA gap is filled in by the action of the DNA polymerase III holoenzyme and the nick is sealed by DNA ligase.

Unlike MMR in *E. coli* and its closely related bacteria, MMR in many other bacteria involves MutL proteins that function as endonucleases (23–26). Studies of the mechanisms of bacterial MutL endonuclease-dependent MMR revealed that some of them rely on MutL endonuclease-β clamp interactions and others do not (24,27).

Eukaryotes contain both MutS and MutL homologs. The MutS homologs MutSα (MSH2-MSH6) and MutSβ (MSH2-MSH3) are sensors of certain mismatches in DNA (28–31). MutSα is the major mismatch recognition factor that recognizes base-base mismatches and small insertion/deletion loops. MutSβ recognizes small insertion/deletion loops. The MutL homolog MutLα (MLH1-PMS2 heterodimer in humans and Mlh1-Pms1 heterodimer in *S. cerevisiae*) acts as an endonuclease in eukaryotic MMR. Germline mutations in the human *MLH1*, *MSH2, MSH6*, and *PMS2* genes cause Lynch and Turcot syndromes (32). Lynch syndrome is characterized by an early onset of colorectal, endometrial, ovarian, and certain other cancers in adults, whereas Turcot syndrome is characterized by brain tumors in children. Lynch syndrome mutations are not evenly distributed in the four MMR genes, with ∼42% in *MLH1*, ∼33% in *MSH2*, ∼18% in *MSH6*, and ∼8% in *PMS2* (33). In addition to Lynch and Turcot syndromes, MMR deficiency causes a subset of sporadic cancers in several human organs (34).

The finding that inactivation of the MMR system by deletion of the *MSH2* gene increases the spontaneous genome-wide mutation rate in *S. cerevisiae* by ∼100-fold emphasizes the critical role of this DNA repair system in maintaining whole-genome stability (35). Eukaryotic MMR proceeds via a series of coordinated events that include mismatch recognition, incision of the daughter strand near the mismatch, and mismatch removal (9,26,36). Biochemical studies have resulted in the reconstitution of MutLα-dependent and independent MMR pathways from purified components (37–39). A critical step in MutLα-dependent MMR pathways is a mismatch-, MutSα-, PCNA-, RFC-, and ATP-dependent incision of the discontinuous strand by the endonuclease activity of MutLα (23). A MutLα-generated strand break located 5’ to the mismatch is utilized by MutSα-activated EXO1 to excise the mismatch in a 5’→3’ hydrolytic reaction modulated by RPA (23,40). The generated gap is filled by the DNA polymerase δ (Pol δ) or Pol χ holoenzyme (37,41,42). When EXO1 is not available, a 5’ strand break created by the activated MutLα endonuclease is utilized by the DNA polymerase δ holoenzyme to remove the mismatch in a strand-displacement DNA synthesis reaction (38,42) enhanced by the nuclease activity of DNA2 (39).

In this study, we investigated MutLα-dependent and independent mechanisms of MMR at a whole genome level in *S. cerevisiae*. We show that MutLα plays a major role in genome-wide MMR in yeast, preferentially defends noncoding DNA against mutations, and suppresses error-prone MMR in yeast and human cells. We also show that a yeast MutLα-independent mechanism contributes to genome-wide MMR.

## RESULTS

### MutLα plays a major role in yeast genome-wide MMR

Yeast MutLα (Mlh1-Pms1 heterodimer) is required for MMR events that take place in the *CAN1*, *his7-2*, and *lys2* mutation reporters (43). However, the involvement of MutLα in MMR events that occur across the whole yeast genome has not been defined. Furthermore, it has not been understood whether a MutLα-independent MMR mechanism promotes genetic stability at a whole-genome level. We initiated this study to examine the contributions of MutLα-dependent and independent mechanisms to genome-wide MMR in *S. cerevisiae*. We first passaged multiple isolates of diploid *pms1Δ*, *msh2Δ*, and wild-type strains for ∼900 generations/isolate to allow them to accumulate *de novo* mutations. We then performed whole genome sequencing and identified *de novo* mutations in each of the passaged isolates by removing variants that were present in the isolate at generation 0 from the list of variants that were present in the same isolate at generation 900. The identified de *novo* mutations that are henceforth referred to as mutations were pooled according to the genotype to generate the mutation spectra (**Table S1**).

To determine the contribution of MutLα to genome-wide MMR, we calculated spontaneous genome-wide mutation rates in the wild-type, *pms1Δ*, and *msh2Δ* strains **(Fig. 1** and **Table S2**). As shown in **Fig. 1A**, the genome-wide mutation rate in the *pms1Δ* strain was 130 times higher than that in the wild-type strain. Importantly, the genome-wide mutation rates in the *pms1Δ* and *msh2Δ* strains did not significantly differ from each other (**Fig. 1A**). Further calculations showed that the rates of deletions in the *pms1Δ* and *msh2Δ* strains were indistinguishable from each other (**Fig. 1B**). Likewise, no significant difference between the rates of base substitutions, or between the rates of insertions, in the two mutant strains was detected. Together, these data indicate that yeast MutLα is required for most of genome-wide MMR events.

**Figure 1.**
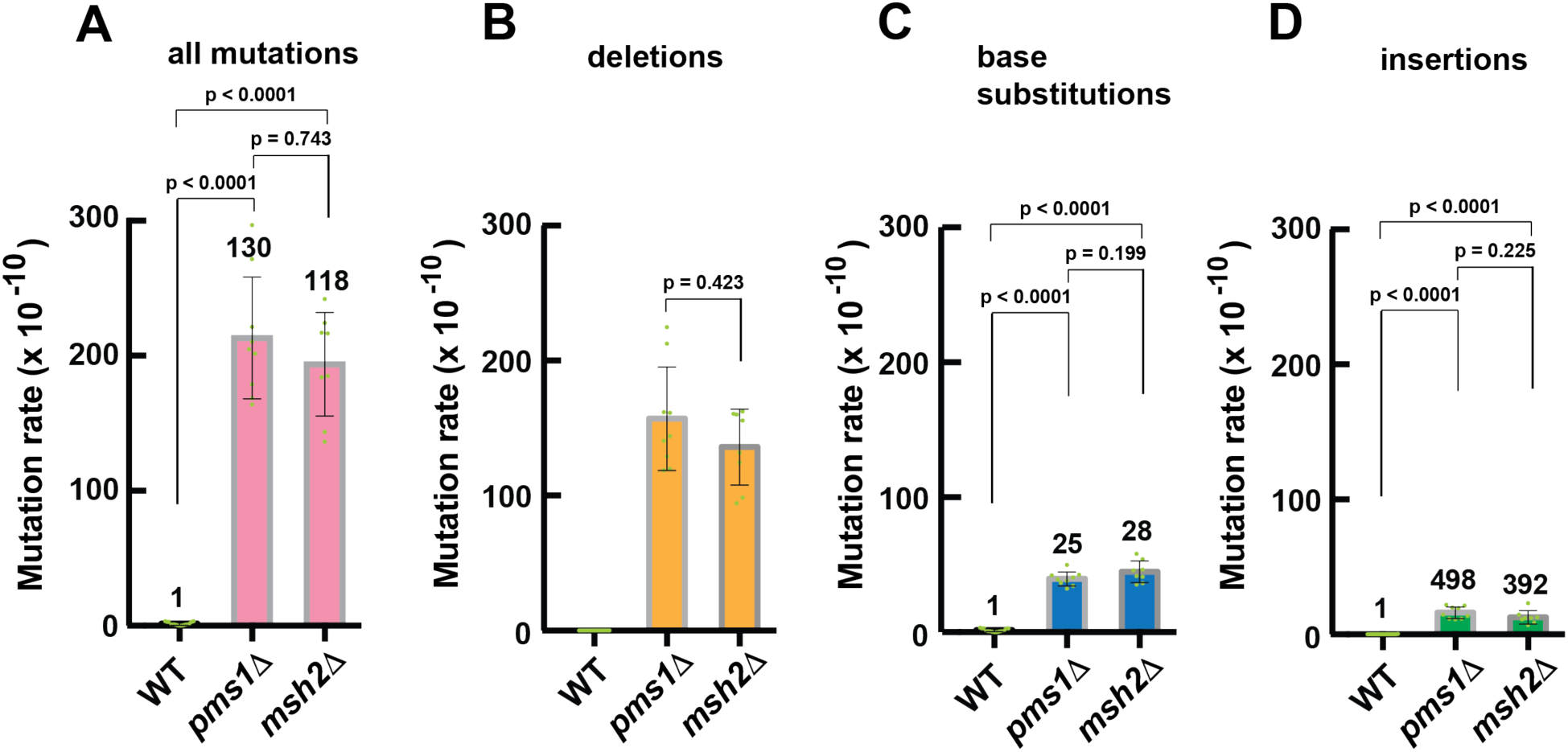
MutLα is required for the majority of MMR events in *S. cerevisiae*. Total rates of genome-wide spontaneous mutations (*A*) and rates of genome-wide spontaneous deletions (*B*), base substitutions (*C*), and insertions (*D*) in the *pms1*Δ, *msh2*Δ, and wild-type strains. The mutation rates were calculated as described in EXPERIMENTAL PROCEDURES and are presented as means ± SD. The data in *A-D* were obtained using independent biological replicates (n_wt_ = 15, n*_pms1_*_Δ_ = 9, and n*_msh2_*_Δ_ = 8). The numbers above the bars are relative mutation rates. The *p* values were calculated using the Mann–Whitney U two-tailed test (GraphPad Prism 6 software). Deletions were not observed in the wild-type strain (*B*).

### Trinucleotide signatures of base substitutions in the *pms1Δ* and *msh2Δ* strains

Analyses of mutational signatures reveal essential information about the nature of mutational processes and DNA repair mechanisms (35,44,45). We extracted trinucleotide signatures of base substitution mutations from the *pms1Δ* and *msh2Δ* strains (**Fig. 2**). The majority of mutations in both signatures are C>A, C>T, and T>C base substitutions. Each of the mutational signatures has three large peaks. Two of the peaks are for C>T base substitutions at the 5’-G<u>C</u>A-3’ and 5’-A<u>C</u>A-3’ sequences, and the third peak is for C>A base substitutions at the 5’-C<u>C</u>T-3’ sequences. The C>T base substitutions at 5’-G<u>C</u>A-3’ and 5’-A<u>C</u>A-3’ sequences and C>A base substitutions at 5’-C<u>C</u>T-3’ sequences account for ∼23% and ∼27% of all base substitutions in the *pms1Δ* and *msh2Δ* strains, respectively. Overall, the two mutational signatures display significant similarities to each other. The observed similarities between the two mutational signatures support the view that MutLα plays a key role in yeast genome-wide MMR.

**Figure 2.**
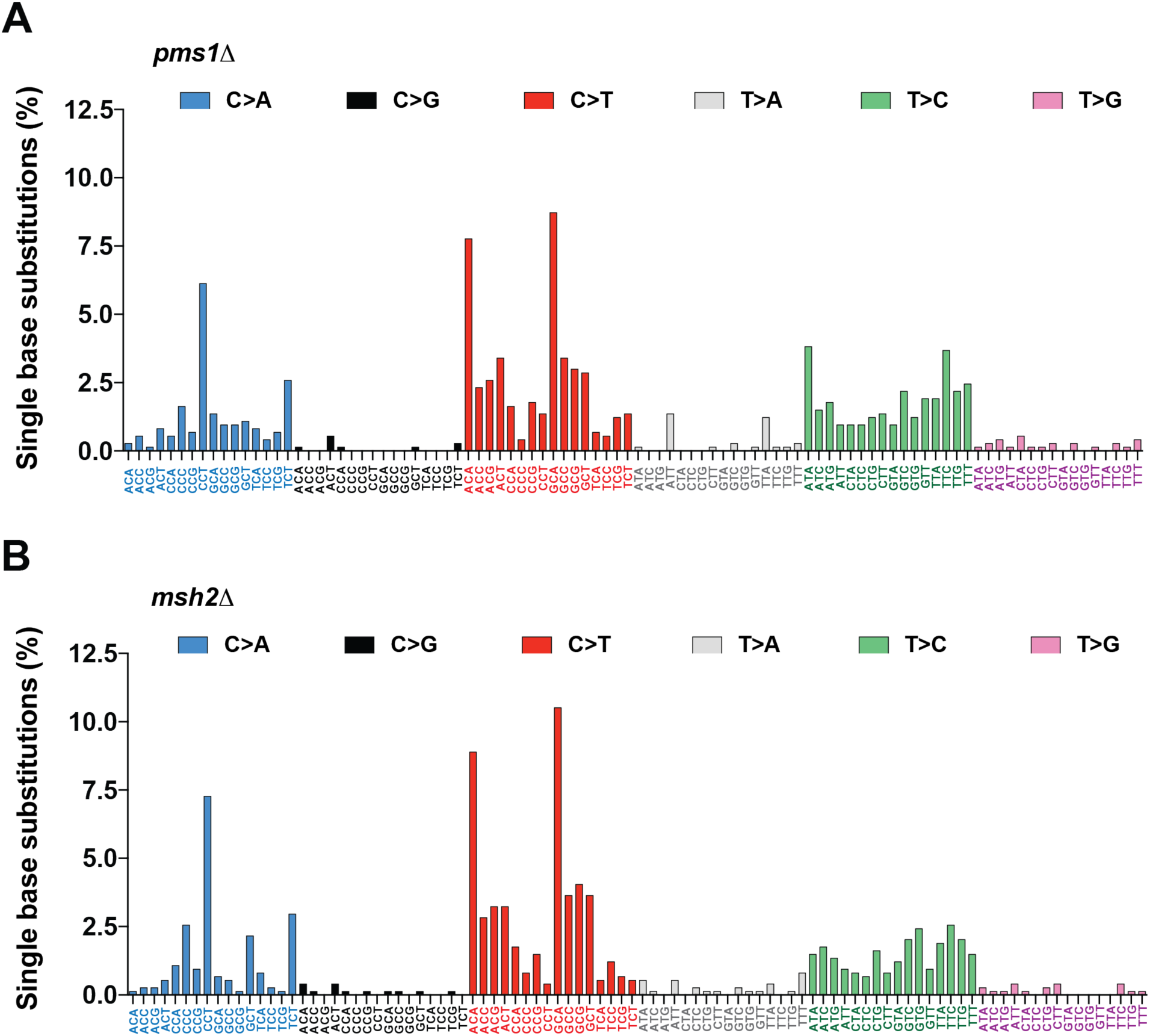
Similarities between trinucleotide signatures of base substitution mutations in MutLα-deficient and Msh2-deficient strains. The trinucleotide mutational signatures were extracted from sequenced nuclear genomes of the *pms1*Δ (*A*) and *msh2*Δ (*B*) strains as described in EXPERIMENTAL PROCEDURES. All base substitution mutations present in the mutation spectra of the *pms1*Δ and *msh2*Δ strains (n*_pms1_*_Δ_ = 734 and n*_msh2_*_Δ_ = 741) were used to generate the trinucleotide mutational signatures.

### MutLα suppresses error-prone MMR in yeast and human cells

It has been unknown whether an MMR factor suppresses an error-prone MMR mechanism that acts at a specific locus or throughout a genome. Our analysis of the trinucleotide base substitution signatures of the *pms1Δ* and *msh2Δ* strains showed that there was a significant difference between the two signatures (**Fig. 2**). Specifically, the base substitution signature of the *pms1Δ* strain contains a peak for T>C substitutions at 5’-A<u>T</u>A-3’ sequences which is significantly larger than the corresponding peak present in the mutational signature of the *msh2Δ* strain (p=0.0147) (**Figs. 2-3**). This finding indicates that yeast MutLα suppresses a genome-wide error-prone MMR mechanism that generates T>C base substitutions at 5’-A<u>T</u>A-3’ sequences.

**Figure 3.**
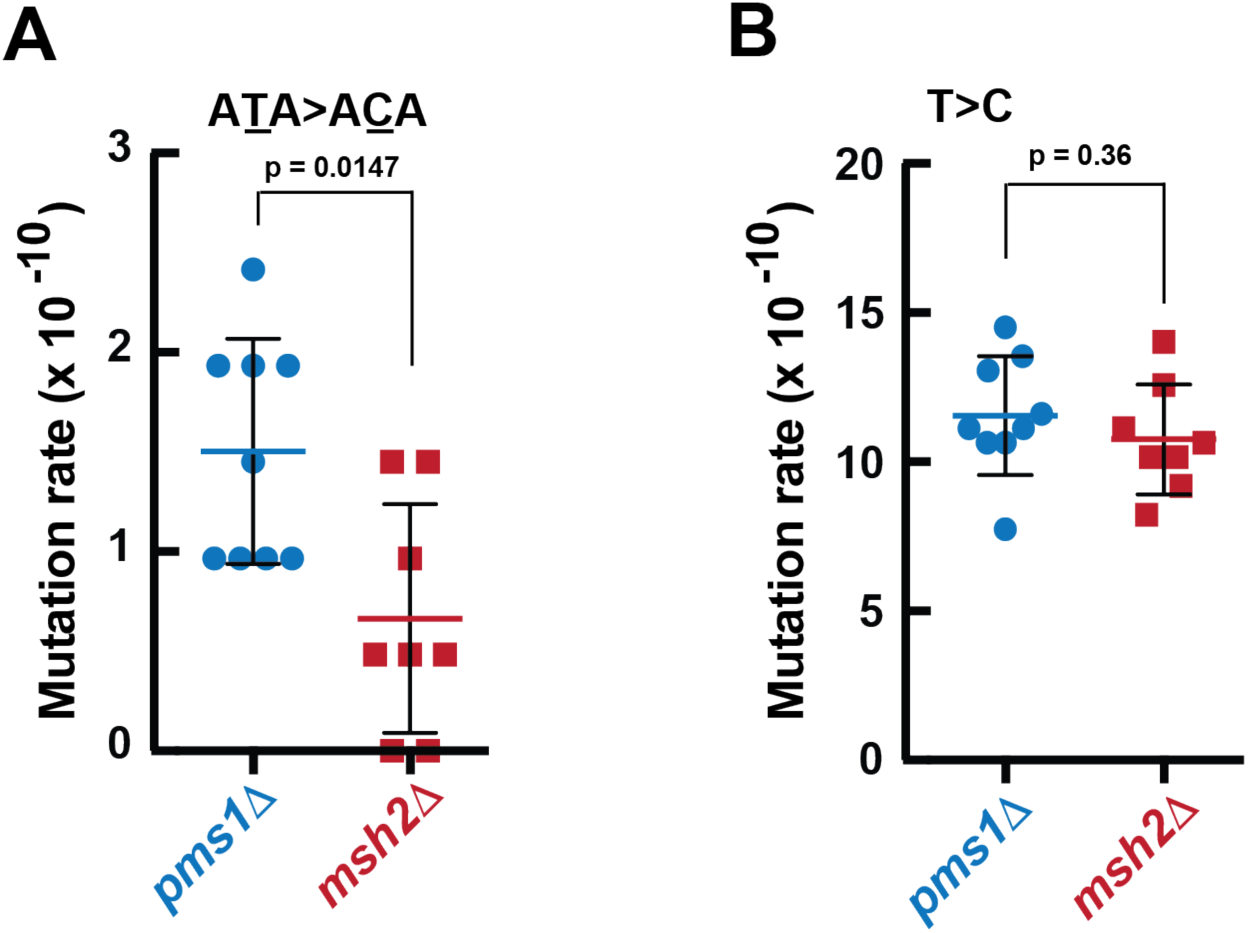
MutLα suppresses error-prone MMR in *S. cerevisiae*. Rates of spontaneous genome-wide T>C base substitutions in 5’-A<u>T</u>A-3’(*A*) and 5’-N<u>T</u>N-3’ (*B*) sequences in the *pms1*Δ and *msh2*Δ strains. The Mann–Whitney U two-tailed test (GraphPad Prism 6 software) was used to determine the *p* values.

To study whether MutLα suppresses error-prone MMR in higher eukaryotes we analyzed recently published mutation data that were obtained by whole genome sequencing of human *ΔMSH2* and *ΔPMS2* induced pluripotent stem cells (iPSCs) (46). The results of our analysis showed that the rate of T>C base substitutions is significantly higher in *ΔPMS2* iPSCs that lack MutLα (MLH1-PMS2 heterodimer) than *ΔMSH2* iPSCs (**Fig.4A**). In contrast, the total rate of base substitutions in *ΔPMS2* iPSCs is significantly lower than the total rate of base substitutions in *ΔMSH2* iPSCs (46). We also observed that one of the peaks of T>C substitutions in the trinucleotide mutational signature of *ΔPMS2* iPSCs is at 5’-A<u>T</u>A-3’ sequences (46) and that the rate of T>C substitutions at 5’-A<u>T</u>A-3’ sequences in the *ΔPMS2* iPSCs is higher than the rate of T>C substitutions in the identical sequences in the *ΔMSH2* iPSCs (**Fig. 4B**). Collectively, these data demonstrate that MutLα suppresses error-prone MMR in yeast cells and human iPSCs on the genome-wide level.

**Figure 4.**
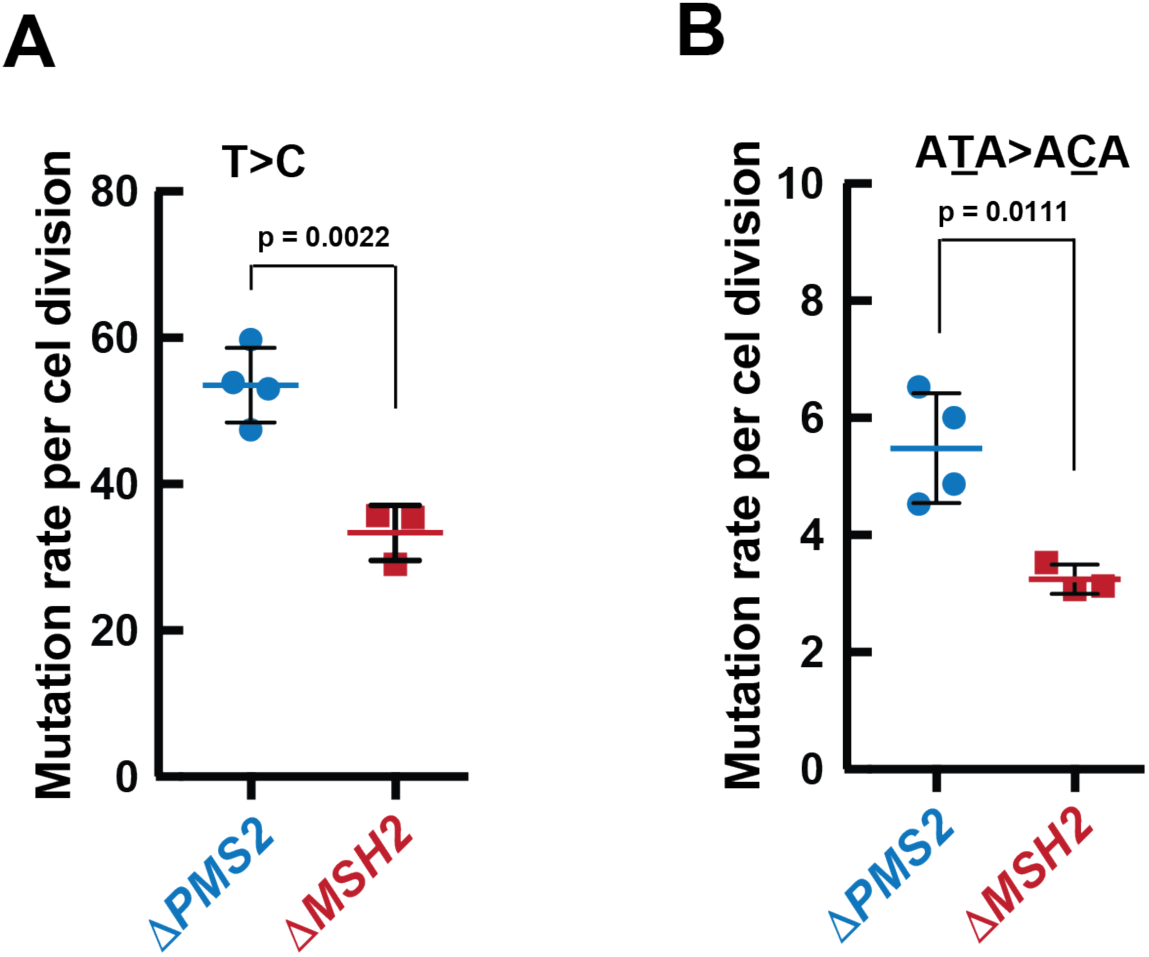
MutLα suppresses error-prone MMR in human iPSCs. (*A*) Genome-wide rates of T>C substitutions in 5’-N<u>T</u>N-3’ (*A*) and 5’-A<u>T</u>A-3’ (*B*) sequences in human Δ*PMS2* and Δ*MSH2* iPSCs. The genome-wide mutation rates were calculated as the number of base substitutions per number of cell divisions and are shown as means ± SD. The *p* values were determined by unpaired t-test (GraphPad Prism 6 software). The genome-wide mutations and the mutated sequences are from a previous study that utilized independent biological replicates (n_Δ_*_PMS2_* = 4; n_Δ_*_MSH2_* = 3) (46).

### MutLα preferentially protects non-coding DNA from mutations

Previous research revealed that microsatellite instability, a hallmark of MMR deficiency, in colorectal and endometrial cancer genomes preferentially occurs in noncoding DNA (47). We next analyzed how mutations were distributed between coding and noncoding DNAs in the wild-type and *pms1Δ* strains. In agreement with a previous study (45), we observed that mutations in the wild-type strain did not show a preference for accumulation in coding or noncoding DNA (**Fig. 5A**). In contrast, mutations in the *pms1Δ* strain preferentially accumulated in noncoding DNA (**Fig. 5A**). In fact, there was an ∼8-fold preference for accumulation of mutations in noncoding DNA in the *pms1Δ* strain (**Fig. 5B**). Further analysis showed that in the MutLα-lacking strain the preferences for accumulation of deletions and insertions in noncoding DNA were 10-fold and 8-fold, respectively (**Fig. 5C-D**). These findings show that MutLα preferentially defends noncoding DNA against mutations.

**Figure 5.**
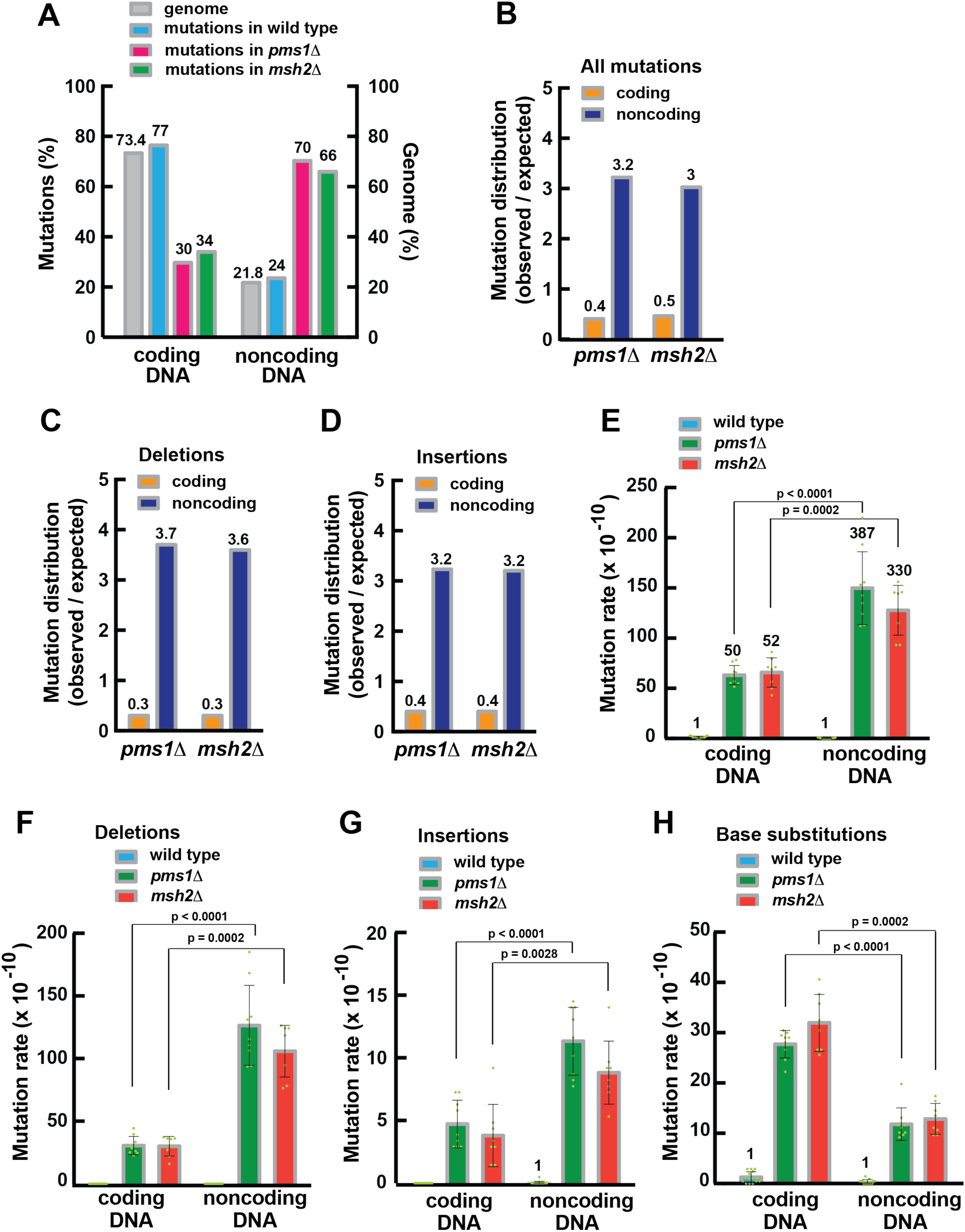
MutLα preferentially defends noncoding DNA against mutations. (*A*) Distribution of genome-wide mutations in coding and noncoding nuclear DNA of the *pms1*Δ, *msh2*Δ, and wild-type strains. (*B*) Genome-wide mutations in coding and noncoding DNA of the *pms1*Δ and *msh2*Δ strains. The numbers above the bars are ratios of the observed to expected mutations. (*C*-*D*) Genome-wide deletions (*C*) and insertions (*D*) in coding and noncoding DNA of the *pms1*Δ and *msh2*Δ strains. The numbers above the bars are ratios of the observed to expected mutations. (*E*) Genome-wide mutation rates in coding and noncoding DNA of the wild-type, *pms1*Δ, and *msh2*Δ strains. The numbers above the bars are relative mutation rates. (*F*-*H*) Genome-wide rates of deletions (*F*), insertions (*G*), and base substitutions (*H*) in coding and noncoding DNA of the wild-type, *pms1*Δ, and *msh2*Δ strains. Data that are shown in *A-H* were obtained using independent biological replicates (n_wt_ = 15, n*_pms1_*_Δ_ = 9, and n*_msh2_*_Δ_ = 8). The *p* values were determined by the Mann–Whitney U two-tailed test (GraphPad Prism 6 software).

A previous study revealed that Msh2 is necessary for preferential protection of noncoding DNA from deletions (35). In line with that study, we determined that Msh2 also preferentially protected noncoding DNA from both insertions (**Fig. 5D**).

### MutLα-independent MMR at a whole genome level

Biochemical studies with purified proteins revealed a MutLα-independent mechanism that corrects mismatches *in vitro* (37,40). However, it has remained unclear whether MutLα-independent MMR functions *in vivo*. To study whether MutLα-independent MMR acts at a whole genome level, we took advantage of Web Logo (48) to visualize patterns in which T>C, C>T, and C>A mutations accumulated in the *pms1Δ* and *msh2Δ* strains. We determined that the sequence pattern in which T>C substitutions accumulated in the *pms1Δ* strain was different from that in the *msh2Δ* strain (**Fig. 6A**). In contrast, the sequence patterns in which C>T or C>A substitutions amassed in the *pms1Δ* and *msh2Δ* strains were similar (**Fig. 6B-C**). A subsequent analysis showed that the most frequent nucleotide that was immediately downstream from A_11_ runs in which 1-bp deletions accumulated was a G in *pms1Δ* cells and a T in *msh2Δ* cells (**Fig. 7**). These analyses indicate that genome-wide MutLα-independent MMR contributes to the suppression of T>C base substitutions and 1-bp deletions in A_11_ runs.

**Figure 6.**
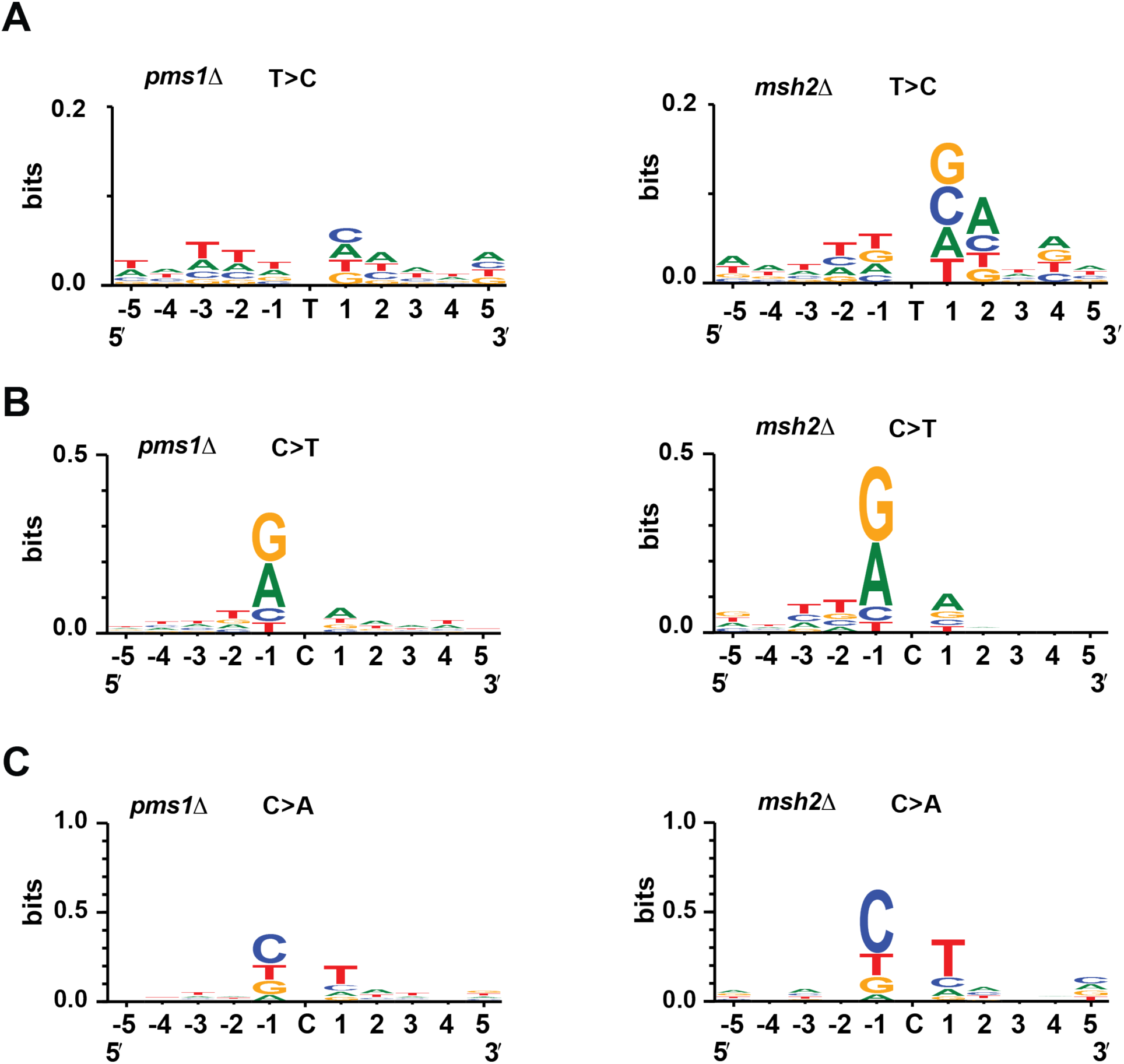
MutLα-independent MMR contributes to the suppression of T>C mutations. Sequence patterns for T>C (*A*), C>T (*B*), and C>A (*C*) base substitutions in the *pms1*Δ and *msh2*Δ strains. The sequence logos were generated using WebLogo 3 as described in EXPERIMENTAL PROCEDURES. To produce the sequence logos, 213 (*A*), 316 (*B*), and 144(*C*) mutated sequences in the *pms1*Δ strain and 177 (*A*), 352 (*B*), and 154(*C*) mutated sequences in the *msh2*Δ strain were analyzed.

**Figure 7.**
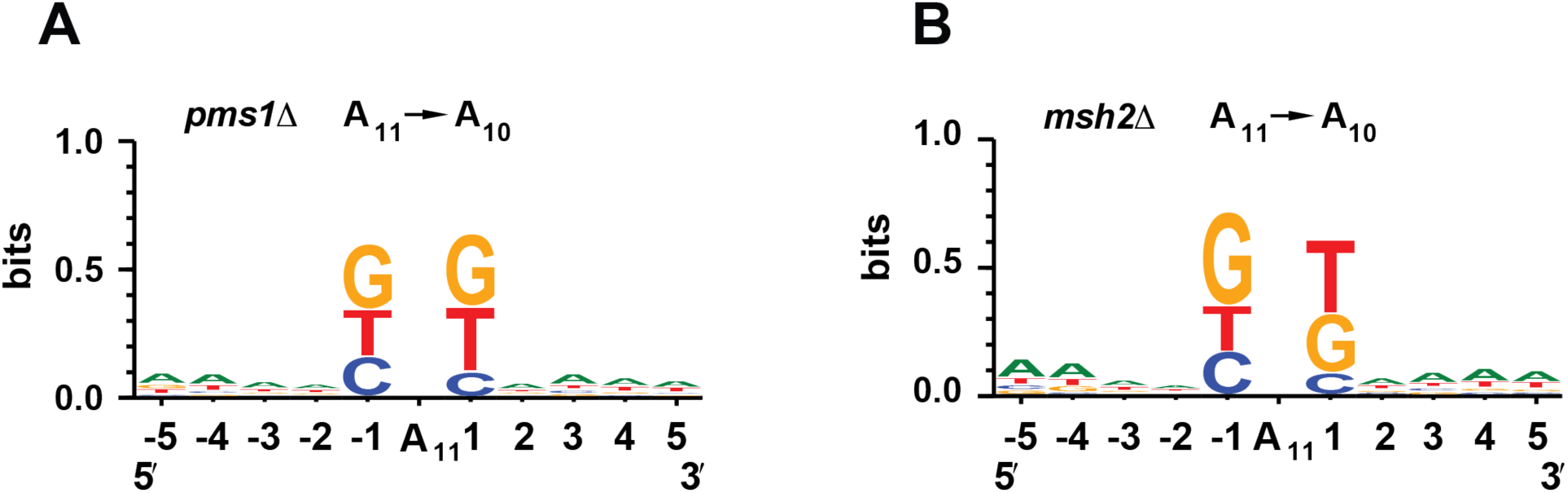
MutLα-independent MMR contributes to the suppression of 1-bp deletions in homopolymeric A_11_ runs. Sequence patterns in which 1-bp deletions accumulated in the *pms1*Δ (*A*) and *msh2*Δ (*B*) strains were obtained with WebLogo 3 as described in EXPERIMENTAL PROCEDURES. The mutated sequences that were analyzed to prepare the sequence logos were 366 (*A*) and 271 (*B*).

Base substitutions in the *pms1Δ* and *msh2Δ* strains frequently occurred in homopolymeric runs (**Fig. 8A**). We next analyzed the involvement of MutLα-independent MMR in the suppression of base substitutions in homopolymeric runs. C>T mutations are the most common base substitutions in the mutation spectra of the *pms1Δ* and *msh2Δ* strains (**Table S1** and **Fig. 2**). Our initial analysis showed that C>T base substitutions in homopolymeric N_3_ runs accumulated at a higher rate in the *msh2Δ* than *pms1Δ* strain (**Fig. 8B**). A following analysis revealed that the total rate of base substitutions in N_3_ runs was higher in the *msh2Δ* strain compared to the *pms1Δ* strain (**Fig. 8C**). Therefore, MutLα-independent MMR is more important for the suppression of base substitutions in homopolymeric N_3_ runs than MutLα-dependent MMR.

**Figure 8.**
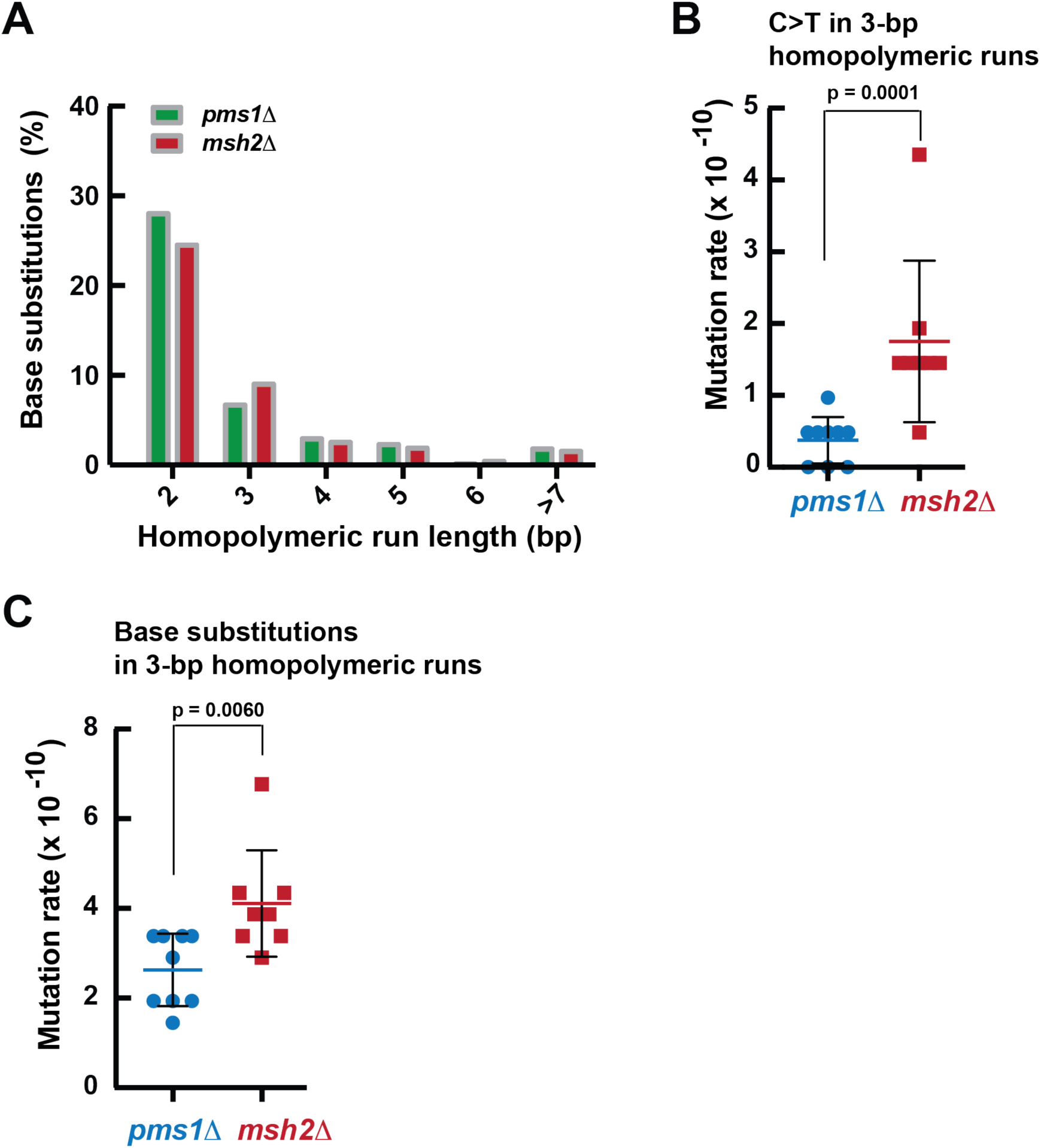
MutLα-independent MMR plays a role in the suppression of base substitutions in N_3_ homopolymeric runs. (*A*) Distribution of base substitution mutations in homopolymeric sequences of different length. (*B*) Rates of C>T transitions in 3-nt homopolymeric runs of the Pms1-and Msh2-deficient strains. (*C*) Rates of base substitutions in 3-nt homopolymeric runs of the *pms1*Δ and *msh2*Δ strains. The mutation rates in *B-C* were calculated as described in the EXPERIMENTAL PROCEDURES and are presented as means ± SD. The Mann–Whitney U two-tailed test (GraphPad Prism 6 software) was used to compute the *p* values.

## Discussion

*E. coli* MutL is the best-studied bacterial MutL protein. It functions as a molecular matchmaker in MMR by coupling mismatch recognition by MutS to the downstream steps of the repair process (1). *E. coli* MutL lacks the DQHA(X)_2_E(X)_4_E endonuclease motif and does not have an endonuclease function in MMR (23,43). The endonuclease function in *E. coli* MMR is provided by MutH. The majority of bacteria and eukaryotes lack MutH endonuclease and its homologs. In these organisms, MutL homologs act as endonucleases in MMR (23–26,43,49,50). MutLα is the founding member of the MutL endonuclease family (23,43). MutLα functions downstream from the mismatch recognition step that is accomplished by MutSα or MutSβ (23,51). In this study, we investigated the involvement of MutLα in genome-wide MMR in *S. cerevisiae*. We have found that loss of MutLα caused by deletion of the *PMS1* gene elevates the genome-wide mutation rate to a level that is present in the *msh2Δ* strain (**Fig. 1A**). Because MMR does not function in the absence of *MSH2*, the finding that the genome-wide mutation rates in the *msh2Δ* and *pms1Δ* strains do not significantly differ from each other (**Fig. 1**) demonstrates that MutLα is required for most of MMR events occurring across the yeast nuclear genome.

Mutational signatures have been instrumental in understanding mutational processes operating in human cancers as well as replication fidelity and DNA repair mechanisms (44,45,52). We extracted and examined the trinucleotide signatures of base substitution mutations from the *pms1Δ* and *msh2Δ* strains (**Fig. 2**). As described below, these two mutational signatures display significant similarities to SBS44, a mutational signature of human MMR deficiency that was extracted from cancers (52). First, the majority of base substitutions in all three mutational signatures are C>T, T>C, and C>A alterations. Second, the three most prominent peaks that are present in the mutational signatures of *PMS1* and *MSH2* deficiency (**Fig. 2**) are also present in SBS44. These similarities provide strong evidence that DNA polymerases in yeast and human cells produce similar base-base mismatches in the same sequence contexts.

Error-prone MMR occurs in variable regions of immunoglobulin genes and certain triplet repeat loci (8,14,15). Because MMR includes the step of DNA re-synthesis it is not surprising that MMR can be error-prone. MutLα has been best known for providing the endonuclease function for error-free MMR. We show that the loss of yeast MutLα triggers error-prone MMR (**Fig. 3**). In line with this finding, our analysis of the human whole genome mutational data demonstrates that MutLα deficiency in the iPSCs also leads to error-prone MMR (**Fig. 4**). Thus, not only does MutLα play a major role in the error-free correction of DNA polymerase errors, but it also suppresses error-prone MMR. The finding that MutLα deficiency results in error-prone MMR in yeast and human cells improves our understanding of the contribution of MutLα to the correction of DNA replication errors. The mechanism of error-prone MMR occurring in response to the absence of MutLα is not known. Because a low-fidelity DNA polymerase, Pol ρι, and EXO1 are involved in error-prone MMR in somatic hypermutation of immunoglobulin genes (8,15), it is possible that these two enzymes also contribute to error-prone MMR that is triggered by the loss of MutLα. It would be important to define this error-prone mechanism using genetic and biochemical approaches.

Noncoding DNA contains gene promoters and other critical regulatory elements. In *S. cerevisiae*, many promoters include homopolymeric dA:dT sequences that are required for normal levels of transcription (53,54). Importantly, the reduction of the size of promoter homopolymeric dA:dT sequences significantly decreases the level of transcription (53). In agreement with this finding, transcription of yeast *HIS3* was stimulated 3-fold after incorporation of a 17-bp poly(dA:dT) sequence in its promoter region (53). Therefore, it is beneficial for the organism to maintain the stability of noncoding DNA. A previous study revealed that Msh2 is required for preferential defense of noncoding DNA against deletions (35). We show here that MutLα preferentially protects noncoding DNA from deletions and insertions but not from base substitutions (**Fig. 5F-H**). Our data analysis indicates that loss of MutLα causes an ∼10-fold bias for the accumulation of deletions and insertions in noncoding relative to coding DNA (**Fig. 5C-D**). Deletion of *MSH2* leads to the formation of the same biases (**Fig. 5C-D**). Of importance is the observation that in the *pms1Δ* and *msh2Δ* strains the mutation rates in noncoding DNA are ∼7 times higher than the mutation rates in coding DNA (**Fig. 5E**), despite the fact that the size of yeast noncoding DNA is ∼3 times smaller than that of yeast coding DNA (55). The increased rates of mutations in noncoding DNA of the *pms1Δ* and *msh2Δ* strains are due to the elevated rates of deletions and insertions, but not base substitutions (**Fig. 5F-H**). The following observations explains the preferential defense of noncoding DNA against indels by MMR. First, relative to coding DNA noncoding DNA has a significantly larger number of longer homopolymeric sequences (56). Second, longer homopolymeric runs are much more unstable than the shorter ones (57). Third, MMR is much more efficient in correcting insertion/deletion loops in longer than shorter homopolymeric runs (57). The more efficient removal of indel loops at longer poly(dA:dT) runs by the MMR system is probably a result of reduced nucleosome density at these homopolymeric runs (58).

Biochemical studies reconstituted a human MutLα-independent MMR pathway that corrects mismatches on 5’-nicked DNA (37,40). This MutLα-independent pathway relies on the mismatch recognition factor MutSα and the 5’→3’ exonuclease activity of EXO1 to excise mismatches in *vitro*. However, the contribution of this pathway to MMR in eukaryotic organisms in *vivo* had remained unclear. We investigated whether yeast MutLα-independent MMR occurred throughout the nuclear genome (**Figs. 6-8**). We observed that the sequence pattern for T>C mutations in the *pms1Δ* cells is different from the sequence pattern for the same mutations in the *msh2Δ* cells (**Fig. 6A**). Likewise, the sequence pattern for 1-bp deletions in homopolymeric A_11_ runs of *pms1Δ* cells differs from the sequence pattern for 1-bp deletions in homopolymeric A_11_ runs of *msh2Δ* cells (**Fig. 7**). Furthermore, we determined that in N_3_ homopolymeric runs the rates for C>T and other base substitutions in the *msh2Δ* cells is significantly higher than the rates for the same genetic alterations in the *pms1Δ* cells. Collectively, these data indicate that MutLα-independent MMR contributes to the maintenance of whole genome stability in *S. cerevisiae*. Assessment of the mutation rates in human *ΔMSH2* and *ΔPMS2* iPSCs supports the view that a MutLα-independent process plays a role in genome-wide MMR (46).

In summary, we show here that MutLα plays a major role in genome-wide MMR, suppresses error-prone MMR, and preferentially defends noncoding DNA against mutations. We also show that MutLα-independent MMR functions *in vivo*.

## EXPERIMENTAL PROCEDURES

### S. cerevisiae strains

Yeast wild-type haploid strains that were used in this study are isogenic BY4741 (*MATa his3Δ1 leu2Δ0 met15Δ0 ura3Δ0*) and BY4742 (*MATα his3Δ1 leu2Δ0 lys2Δ0 ura3Δ0*), both of which are derivatives of S288C. *PMS1* and *MSH2* gene deletions in the haploid wild-type strains were generated by lithium/PEG-based transformations of PCR-amplified gene replacement cassettes. The gene deletions were confirmed by PCRs. Homozygous diploid yeast strains lacking *PMS1* (FKY2291, FKY2292, FKY2293, and FKY2294) or *MSH2* (FKY1982, FKY1983, and FKY1984) were constructed by crossing haploid BY4741 and BY4742 strains carrying *pms1Δ* or *msh2Δ*. Wild-type diploid strains FKY1719, FKY1720 and FKY1721 are isogenic to the homozygous diploid *pms1Δ* or *msh2Δ* strains and were described previously (45).

### Mutation accumulation, library preparation, and genome sequencing

To amass spontaneous mutations in the diploid yeast *pms1Δ*, *msh2Δ*, and wild-type strains mutation accumulation experiments were performed. In these mutation accumulation experiments, we utilized 30 single-cell bottlenecks to passage fifteen wild-type, nine *pms1Δ,* and eight *msh2Δ* isolates for ∼900 generations at 30°C on solid YPDAU medium (45). Glycerol stocks of the passaged isolates that were at generations 0 and 900 were prepared and stored at −80°C.

The glycerol stocks were used to prepare patches of the multiple isolates of the passaged yeast *pms1Δ*, *msh2Δ*, and wild-type strains on solid YPDAU medium at 30°C for 20-24 hours. Genomic DNAs from the fresh patches were isolated using a MasterPure DNA purification kit (LGC Bioresearch Technologies). 400 ng genomic DNA of each sample was used to construct whole-genome DNA libraries with an NEBNext Ultra II FS DNA Library prep kit (NEB) and NEBNext Multiplex Oligos for Illumina (NEB). DNA fragments were size-selected for an average insert size of 450 bp. The libraries were analyzed using a TapeStation system (Agilent). The 151-bp 2D paired-end sequencing was performed using a NovaSeq 6000 sequencing system (Illumina) with an S4 PE 2×150 flow cell in XP mode.

### Mutation spectra and calculation of mutation rates

The initial Binary Base Calls (BCL) sequencing files were demultiplexed and converted to FASTQ files using bcl2fastq Conversion Software, v. 2.20.0 (Illumina). The obtained paired-end sequencing data were imported into CLC Genomics Workbench (Qiagen) and aligned to the *S. cerevisiae* S288C reference genome. Mutations were called as described previously (45). Variants present within repetitive elements were not called if they could not be uniquely mapped. Mutations that were present in an isolate at generation 0 were removed from the list of mutations that were present in the same isolate at generation 900. The mutation spectra were generated by pooling mutations accumulated in the multiple isolates of the wild-type, *pms1Δ,* and *msh2Δ* strains according to the genotype.

Mutation rates (µ) were calculated as previously described (45) using the following equation: *µ = N_i_ / gen / N_g_,* where *N_i_* is the number of mutations of type *i*, *N_g_* is the size of the diploid genome in which the variants were called (22,983,805 bp), and *gen* is the total number of generations for all isolates of the genotype.

### Mutational signatures and sequence logos

The trinucleotide signatures of base substitution mutations (44) and the sequence logos were generated as previously described (45). Briefly, the position of each mutation in the genome and the 5′ and 3′ flanking sequences were determined using the *S. cerevisiae* S288C reference genome sequence and the CLC Genomics Workbench (Qiagen). The mutated trinucleotide sequences were sorted into the 96 different classes (44) using the Excel Data Filter Tool (Microsoft). WebLogo 3 (48) (https://weblogo.threeplusone.com/create.cgi) was used to prepare sequence logos.

## DATA AVAILABILITY

Whole genome DNA sequencing data will be available in the Sequence Read Archive database at the National Center for Biotechnology Information at the time of publication.

## SUPPORTING INFORMATION

This article contains supporting information.

## ACKNOWLEDGEMENTS

We thank the High Throughput Sequencing Facility at the University of North Carolina at Chapel Hill for sequencing of our whole genome libraries and demultiplexing of the NGS data. We also thank Farid F. Kadyrov for critical reading of the manuscript.

## AUTHOR CONTRIBUTIONS

L.K., P.M., and F.K. designed research; L.K., P.M., and F.K. performed research; L.K., P.M., and F.K. analyzed data; and L.K., P.M., and F.K. wrote the manuscript.

## FOOTNOTES

Research described in this publication was supported by the National Institute of General Medical Sciences of the National Institutes of Health under Award Number R01GM132128 (to F.A.K.). The content is solely responsibility of the authors and does not necessarily represent the official views of the National Institutes of Health.

## CONFLICT OF INTEREST

The authors declare that they have no conflicts of interest with the contents of this article.

## Notes

### Competing Interest Statement

The authors have declared no competing interest.

