## Supplemental Information for "MutLα suppresses error-prone DNA mismatch repair and preferentially protects noncoding DNA from mutations"

Running title: MutLα and genome-wide MMR

MATERIALS INCLUDED:

TABLES S1-S2

**Table S1. Mutation spectra in the wild type, *pms1Δ*, and *msh2Δ* diploid yeast strains**

| | wild type | *pms1Δ* | *msh2Δ* |
|---|---|---|---|
| **Isolates** | 15 | 9 | 8 |
| **Passages** | 450 | 270 | 240 |
| **Generations** | 13,500 | 8,100 | 7,200 |
| **Genome size** | 22,983,805 bp | | |
| **Mutation type** | **Number of mutations in each category (n)** | | |
| **Deletions of single A/T pairs** | 0 | 2,008 | 1,572 |
| **Deletions of single G/C pairs** | 0 | 73 | 68 |
| **Insertions of single A/T pairs** | 1 | 217 | 154 |
| **Insertions of single G/C pairs** | 0 | 17 | 13 |
| **>1-bp deletions** | 0 | 843 | 611 |
| **>1-bp insertions** | 0 | 65 | 42 |
| **T→C** | 6 | 213 | 177 |
| **C→T** | 10 | 316 | 352 |
| **T→A** | 3 | 28 | 27 |
| **T→G** | 5 | 24 | 18 |
| **C→A** | 19 | 144 | 154 |
| **C→G** | 6 | 9 | 13 |
| **other** | 1 | 4 | 3 |
| **Total** | 51 | 3961 | 3,204 |

**Table S2. Mutation rates in the *pms1Δ*, *msh2Δ*, and wild-type diploid yeast strains**

| | wild type | *pms1Δ* | *msh2Δ* |
|---|---|---|---|
| **Isolates** | 15 | 9 | 8 |
| **Passages** | 450 | 270 | 240 |
| **Generations** | 13,500 | 8,100 | 7,200 |
| **Genome size** | 22,983,805 bp | | |
| **Mutation type** | **Absolute rate in each category (x $10^{-11}$)** | | |
| **Deletions of single A/T pairs** | < 0.32 | 1,078 | 950 |
| **Deletions of single G/C pairs** | < 0.32 | 39 | 41 |
| **Insertions of single A/T pairs** | 0.32 | 117 | 93 |
| **Insertions of single G/C pairs** | < 0.32 | 9 | 8 |
| **>1-bp deletions** | < 0.32 | 453 | 369 |
| **>1-bp insertions** | < 0.32 | 35 | 25 |
| **T→C** | 1.9 | 115 | 108 |
| **C→T** | 3.2 | 170 | 213 |
| **T→A** | 0.97 | 15 | 16 |
| **T→G** | 1.6 | 13 | 11 |
| **C→A** | 6.1 | 77 | 92 |
| **C→G** | 1.9 | 4.8 | 8 |
| **other** | 0.32 | 2.1 | 1.8 |
| **Total** | 16.4 | 2,129 | 1,937 |
